# Genome-wide analysis of in vivo CcpA binding with and without its key co-factor HPr in the major human pathogen group A *Streptococcus*

**DOI:** 10.1101/2020.08.28.272682

**Authors:** Sruti DebRoy, Victor Aliaga Tobar, Gabriel Galvez, Srishtee Arora, Xiaowen Liang, Nicola Horstmann, Vinicius Maracaja-Coutinho, Mauricio Latorre, Magnus Hook, Anthony R. Flores, Samuel A. Shelburne

**Affiliations:** Department of Infectious Diseases Infection Control and Employee Health, University of Texas MD Anderson Cancer Center, Houston TX, USA; Department of Genomic Medicine, University of Texas MD Anderson Cancer Center, Houston TX, USA; Facultad de Ciencias Químicas y Farmacéuticas, Advanced Center for Chronic Diseases-ACCDiS, Universidad de Chile,Santos Dumont 964, Independencia, Santiago,Chile; Laboratorio de Bioingeniería, Instituto de Ciencias de la Ingeniería, Universidad de O’Higgins, Libertador Bernardo O’Higgins 611, Rancagua, O’Higgins, Chile; Center for Infectious and Inflammatory Diseases, Institute of Biosciences and Technology, Texas A&M University Health Science Center, Houston, TX, USA; Laboratorio de Bioinformática y Expresión Génica, INTA, Universidad de Chile, El Líbano 5524, Macul, Santiago, Chile; Mathomics, Center for Mathematical Modeling, Universidad de Chile, Beauchef 851, 7th Floor, Santiago, Santiago, Chile; Center for Genome Regulation (Fondap 15090007), Universidad de Chile, Blanco Encalada 2085, Santiago, Santiago, Chile; Centro de Modelamiento Molecular, Biofísica y Bioinformática (CM2B2), Facultad de Ciencias Químicas y Farmacéuticas, Universidad de Chile, Santiago, Chile; Division of Infectious Diseases, Department of Pediatrics, University of Texas Health Science Center McGovern Medical School, Houston, TX, USA; Center for Antimicrobial Resistance and Microbial Genomics, University of Texas Health Science Center McGovern Medical School, Houston, TX, USA

**Keywords:** *Streptococcus pyogenes*, ChIPseq, HPr-independent CcpA regulation

## Abstract

Catabolite control protein A (CcpA) is a master regulator of carbon source utilization and contributes to the virulence of numerous medically important Gram-positive bacteria. Most functional assessments of CcpA, including interaction with its key co-factor HPr, have been performed in non-pathogenic bacteria. In this study we aimed to identify the in vivo DNA binding profile of CcpA and assess the extent to which HPr is required for CcpA-mediated regulation and DNA binding in the major human pathogen group A *Streptococcus* (GAS). Using a combination RNAseq/ChIPseq approach, we found that CcpA affects transcript levels of 514 of 1667 GAS genes (31%) whereas direct DNA binding was identified for 105 GAS genes. Three of the directly regulated genes encode the key GAS virulence factors Streptolysin S, PrtS (IL-8 degrading proteinase), and SpeB (cysteine protease). Mutating CcpA Val301 to Ala (strain 2221-CcpA-V301A) abolished interaction between CcpA and HPr and impacted the transcript levels of 205 genes (40%) in the total CcpA regulon. By ChIPseq analysis, CcpAV301A bound to DNA from 74% of genes bound by wild-type CcpA, but generally with lower affinity. These data delineate the direct CcpA regulon and clarify the HPr-dependent and independent activities of CcpA in a key pathogenic bacterium.

**Data sharing and data availability:** The data that support the findings of this study are available from the corresponding author upon reasonable request.

## Introduction

Carbon catabolite repression (CCR) is a global process by which bacteria prioritize the use of favorable energy sources (Deutscher *et al*., 2006, Fujita, 2009). The core mechanism of CCR is an alteration in levels of proteins involved in metabolite transport and utilization which in turn is primarily achieved at the transcriptional level (Gorke & Stulke, 2008). The LacI-GalR family transcriptional regulator catabolite control protein (CcpA) is a key mediator of CCR in many Gram-positive bacteria (Henkin *et al*., 1991, Titgemeyer & Hillen, 2002, Warner & Lolkema, 2003). CcpA inactivation impacts ∼15-20% of the transcriptome of a broad array of Gram-positive bacteria with the majority of impacted genes encoding proteins involved in carbohydrate and nitrogen utilization (Antunes *et al*., 2012, DebRoy *et al*., 2016, Seidl *et al*., 2009, Zeng *et al*., 2013). Importantly, CcpA also affects the transcript levels of genes encoding known and putative virulence factors in human Gram-positive pathogens ranging from streptococci to *Clostridia* (Iyer *et al*., 2005, Mertins *et al*., 2007, Seidl *et al*., 2008a, Varga *et al*., 2004). Consequently, CcpA inactivation in diverse organisms result in altered virulence-related phenotypes such as extracellular capsule production, biofilm formation, lysis of red blood cells, and inter-species signaling (Giammarinaro & Paton, 2002, Johnson *et al*., 2009, Kinkel & McIver, 2008, Seidl *et al*., 2008b, Watson *et al*., 2013).

Although CcpA is clearly critical to the virulence of numerous key Gram-positive pathogens, understanding of CcpA physiologic function is mainly derived from studies in non-pathogenic bacteria (Warner & Lolkema, 2003). Based primarily on investigations in *Bacillus* species, CcpA is currently thought to impact gene expression by binding *cis*-acting DNA known as catabolite response elements (*cre*) which are composed of the pseudo-palindromic motif WTGNAANCGNWNNCWW (where W = A or T and N = any base) (Miwa *et al*., 2000, Schumacher *et al*., 2011, Stulke & Hillen, 2000). CcpA affinity for *cre* sites is significantly increased by the co-effector molecule, histidine-containing protein (HPr) phosphorylated at Ser46 (HPrSer46∼P) (Aung-Hilbrich *et al*., 2002, Deutscher *et al*., 2005, Shelburne *et al*., 2008). HPr is phosphorylated and dephosphorylated at Ser46 by HPr <u>k</u>inase/<u>p</u>hosphorylase (HPrK/P), a bifunctional ATP-dependent enzyme whose activity is responsive to intracellular energy status (Poncet *et al*., 2004). Thus, the interaction of CcpA with HPr facilitates alteration in gene expression in response to metabolic changes (Deutscher *et al*., 2006). All major, invasive Gram-positive pathogens contain highly conserved orthologues of CcpA, HPr, and HPrK/P with amino acid similarities ranging from 70% for CcpA to 85% for HPr. However, *Bacillus* species also contain Crh (catabolite repression HPr), which is an HPr-like protein important for CcpA-mediated gene regulation that is not present in typical Gram-positive pathogens such as staphylococci and streptococci (Deutscher *et al*., 2006, Galinier *et al*., 1997, Schumacher *et al*., 2006).

Although the pseudo-palindromic *cre* sequence seems to be the predominate site of CcpA-DNA interaction in *Bacillus* species, there have been suggestions that CcpA may bind other DNA sites. Using in silico approaches, *cre* sites have been predicted in only a small percentage of genes whose transcript levels are significantly altered by CcpA inactivation (Carvalho *et al*., 2011, DebRoy *et al*., 2016).

Additionally, a recent study in *Clostridium acetobutylicum* identified a CcpA binding motif (TGTAA/TTTACA) which is quite distinct from the previous *cre* motif (Yang *et al*., 2017). Moreover, in addition to the “classic” *cre* motif, a genome-wide CcpA binding study identified a *cre*2 motif in *Streptococcus suis* (TTTTYHWDHHWWTTTY, where Y is C or T, H is A or C or T, and D is A or G or T) that was primarily important for CcpA function in the stationary phase (Willenborg *et al*., 2014). Although CcpA transcriptome analyses have been performed in a wide variety of bacteria, genome-wide characterization of CcpA-DNA binding is not currently available for a major, invasive Gram-positive pathogen other than *Streptococcus suis* (Antunes *et al*., 2012, Buescher *et al*., 2012, Willenborg *et al*., 2014).

In addition to a sub-optimal understanding of how CcpA interacts with DNA in invasive Gram-positive bacteria, analysis of the role of HPr and HPrK/P in pathogenic bacteria has been quite limited (Mertins *et al*., 2007, Shelburne *et al*., 2008). In part, this is because in major Gram-positive pathogens such as group A *Streptococcus* (GAS), *Streptococcus pneumoniae*, and *Staphylococcus aureus*, HPr or even HPrSer46∼P appears to be essential, in contrast to what is observed in *Bacillus* species (Fleming *et al*., 2015). In *Staphylococus xylosu*s elimination of HPrSer46∼P completely abolished CCR suggesting that CcpA absolutely requires HPrSer46∼P to impact gene transcription (Jankovic & Bruckner, 2002). In contrast, CcpA regulation at *cre*2 sites in *S. suis* was postulated to be independent of HPr (Willenborg *et al*., 2014). As pathogens establish and propagate human infections, they are likely to encounter vastly different metabolic conditions, which in turn would be expected to alter HPrSer46∼P levels. Thus establishing whether CcpA can function independently of HPrSer46∼P and which genes are regulated by CcpA in the absence of HPrSer46∼P is highly pertinent to fully understand the role that nutritional acquisition plays in modulation of pathogenesis and infection outcomes. Herein, we sought to determine the global DNA binding characteristics of CcpA in GAS and analyze the effect of blocking the interaction between CcpA and HPrSer46∼P in order to broaden insight into the physiology underlying the critical contribution of CcpA to Gram-positive pathogenesis.

## Results

### A CcpA mutant that does not bind HPr

We chose to conduct our study in the *emm1* GAS strain MGAS2221 because *emm1* strains are leading causes of GAS infections, MGAS2221 is representative of the current *emm1* strains causing human disease worldwide, and MGAS2221 has a fully sequenced genome and lacks mutations in known regulators that have been shown to impact the CcpA transcriptome (Shelburne *et al*., 2010, Sumby *et al*., 2005). Both HPr and HPrK/P are essential in GAS (Le Breton *et al*., 2015). Thus, to probe into the possible existence of a CcpA regulon independent of HPrSer46∼P, we first attempted to generate an HPr-S46A mutant. However, we were unable to generate a viable GAS HPr-S46A mutant despite repeated efforts suggesting that, as in the case of *Streptococcus pneumoniae* (Fleming *et al*., 2015), a S46A mutation in GAS HPr is lethal, possibly due HPrSer46∼P dependent cellular functions that are essential. As an alternate approach we sought to generate a GAS mutant which retains the ability for HPr to be phosphorylated, but does not permit interactions between HPrSer46∼P and CcpA. In silico alanine scanning mutagenesis of the CcpA-HPrSer46∼P interface, modeled on the crystal structure of the *Bacillus megaterium* CcpA-HPrSer46∼P-DNA complex, identified four amino acid residues of CcpA that are critical to CcpA-HPrSer46∼P interaction (Homeyer *et al*., 2007, Schumacher *et al*., 2004). We chose to alter one of these residues, valine 301, to alanine given that the V301 residue is conserved in all CcpA family proteins (Schumacher *et al*., 2004) and gene regulation studies have demonstrated that a CcpAV301 mutant is compromised in glucose-mediated gene repression (Sprehe *et al*., 2007).

We first sought to test the hypothesis that the CcpAV301A mutation would impair the interaction of recombinant CcpA and HPr using surface plasmon resonance (SPR). To this end, we expressed and purified recombinant CcpA and CcpAV301A along with HPr. We used recombinant HPrK/P to phosphorylate HPr at Ser46 as previously described (Shelburne *et al*., 2010). As expected, wild type CcpA had about 30 fold higher affinity (i.e. bound more tightly) for HPrSer46∼P relative to HPr with *K*_D_ value of 5.8 ± 0.5 vs. 188 ± 18 µM (Figure 1). Consistent with our hypothesis that V301 is critical for CcpA-HPrSer46∼P interaction in GAS, recombinant CcpAV301A bound to HPrSer46∼P with a *K*_D_ value of 153 ± 17 µM which closely approximated that observed for wild type CcpA/HPr interaction (*K*_D_ = 188 ± 18 µM)(Figure 1). The interaction between CcpAV301A and HPr is much weaker and the affinity could not be determined under the conditions used. The equilibrium dissociation constant values (*K*_D_) are mean ± standard deviation obtained from two experiments.

**Figure 1.**
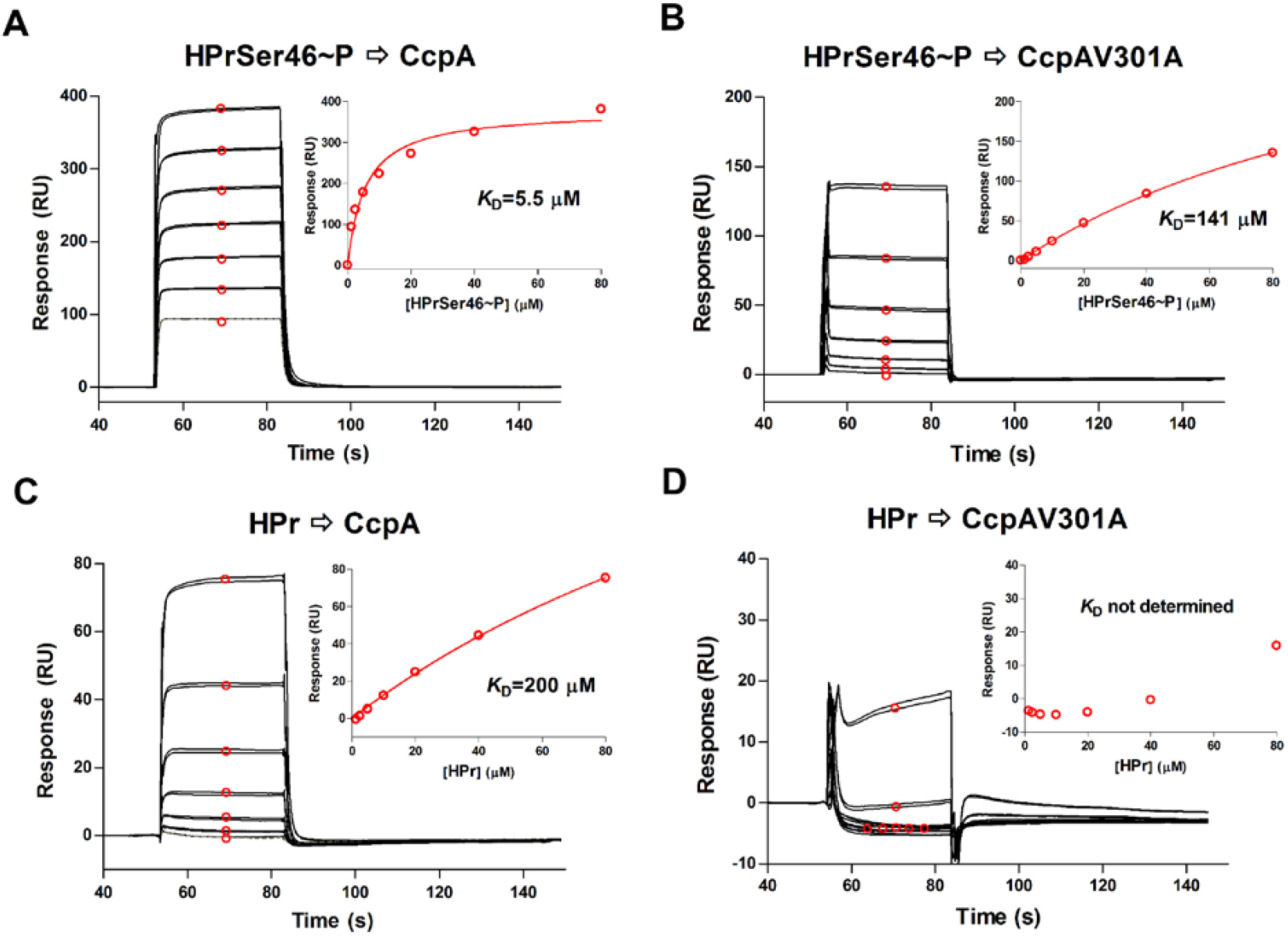
In vitro analysis of CcpA-HPr interaction. Representative SPR analysis (n=2) of the binding between CcpA and HPr recombinant proteins. HPrSer46∼P and HPr (2-fold serial dilutions from 1.25 to 80 μM) was injected in duplicate to (A & C) CcpA surface (3700 RU) and (B & D) CcpAV301A surface (4200 RU). The SPR response curves of bound protein are shown in black with lower curve corresponding to lower concentration of protein injected. The average responses at steady state (shown in red circles) were plotted as a function of the HPr concentration and the isotherm was fit to a one-site binding (hyperbola) model (fitted curve shown in red) to determine equilibrium dissociation constant *K*_D_ (inset).

Next, we used site directed mutagenesis to create the isoallellic mutant strain 2221-CcpA-V301A in the same parental background, the serotype *emm1* strain MGAS2221, as our previously created 2221Δ*ccpA* isolate (Table 1). Given that deleting *ccpA* has previously been shown to affect HPr/HPr∼P ratios in other bacteria (Leboeuf *et al*., 2000, Ludwig *et al*., 2002), we used Phos-tag gels to analyze HPr and HPr∼P levels at the mid-exponential phase of growth in a nutrient rich media. Under these conditions, the majority of HPr in strains MGAS2221 and 2221-CcpA-V301A was unphosphorylated with no significant difference of HPr/HPr∼P ratios identified between the two strains (Figure 2A, B). Conversely, HPr∼P levels were increased in strain 2221Δ*ccpA* relative to the other two strains. A similar increase in HPr∼P levels has been observed in *ccpA* mutants of *B. subtilis* and *E. faecalis*, and has been attributed to in*cre*ased HPr kinase activity (Leboeuf *et al*., 2000, Ludwig *et al*., 2002). These data show that the V301A alteration in CcpA did not have significant effects on cellular HPr∼P levels (Figure 2A, B). Additionally, the cellular levels of CcpA were similar between MGAS2221 and 2221-CcpA-V301A indicating that the amino acid variation did not impact CcpA autoregulation (note similar CcpA band densities for MGAS2221 and 2221-CcpA-V301A in Figure 2C, third panel).

**Table 1.**
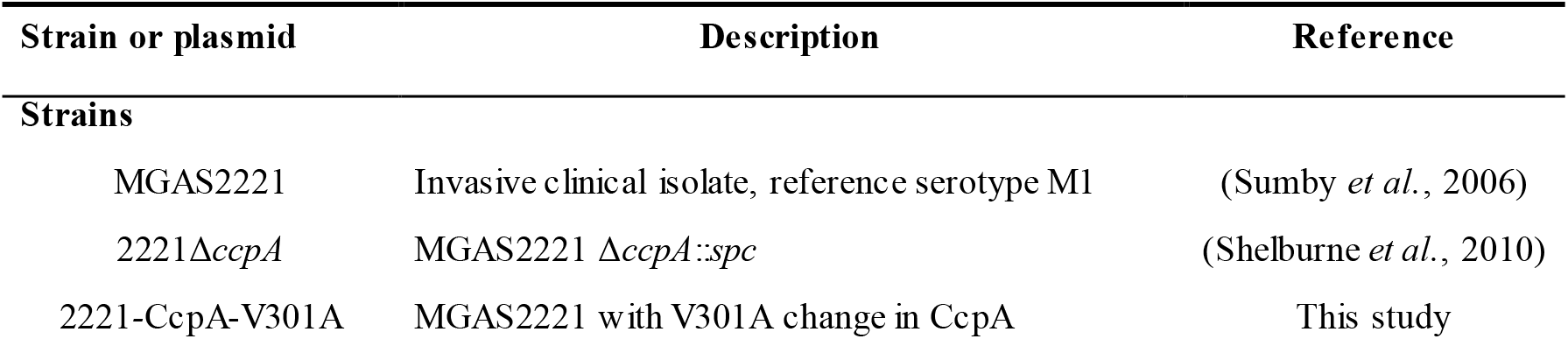
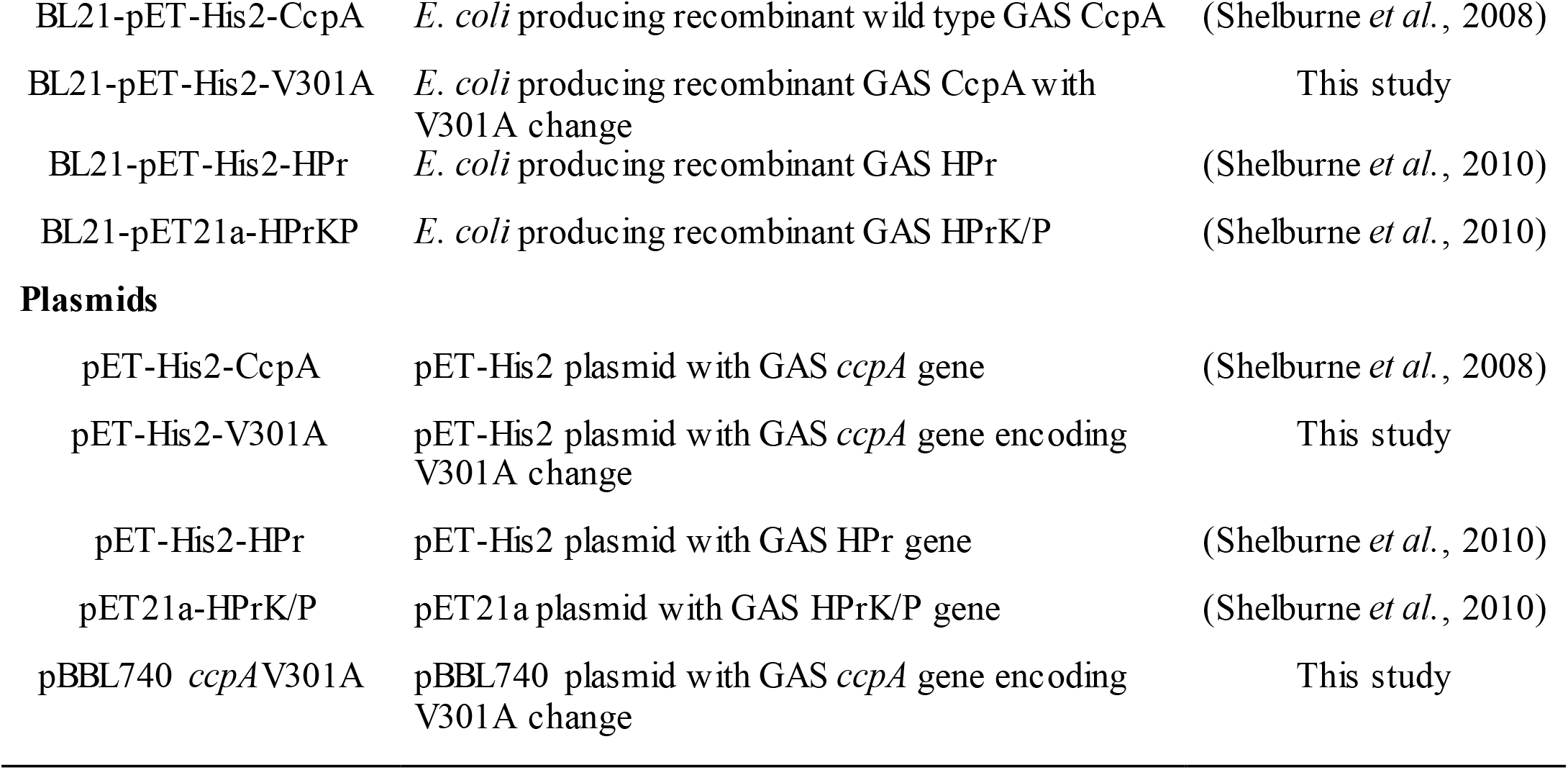
Strains and plasmids used in this study.

| Strain or plasmid | Description | Reference |
| --- | --- | --- |
| **Strains** |  |  |
| MGAS2221 | Invasive clinical isolate, reference serotype M1 | (Sumby *et al.*, 2006) |
| 2221Δ*ccpA* | MGAS2221 Δ*ccpA*::*spc* | (Shelburne *et al.*, 2010) |
| 2221-CcpA-V301A | MGAS2221 with V301A change in CcpA | This study |
| BL21-pET-His2-CcpA | *E. coli* producing recombinant wild type GAS CcpA | (Shelburne *et al.*, 2008) |
| BL21-pET-His2-V301A | *E. coli* producing recombinant GAS CcpA with V301A change | This study |
| BL21-pET-His2-HPr | *E. coli* producing recombinant GAS HPr | (Shelburne *et al.*, 2010) |
| BL21-pET21a-HPrK/P | *E. coli* producing recombinant GAS HPrK/P | (Shelburne *et al.*, 2010) |
| Plasmids |  |  |
| pET-His2-CcpA | pET-His2 plasmid with GAS *ccpA* gene | (Shelburne *et al.*, 2008) |
| pET-His2-V301A | pET-His2 plasmid with GAS *ccpA* gene encoding V301A change | This study |
| pET-His2-HPr | pET-His2 plasmid with GAS HPr gene | (Shelburne *et al.*, 2010) |
| pET21a-HPrK/P | pET21a plasmid with GAS HPrK/P gene | (Shelburne *et al.*, 2010) |
| pBBL740 *ccpAV301A* | pBBL740 plasmid with GAS *ccpA* gene encoding V301A change | This study |

**Figure 2.**
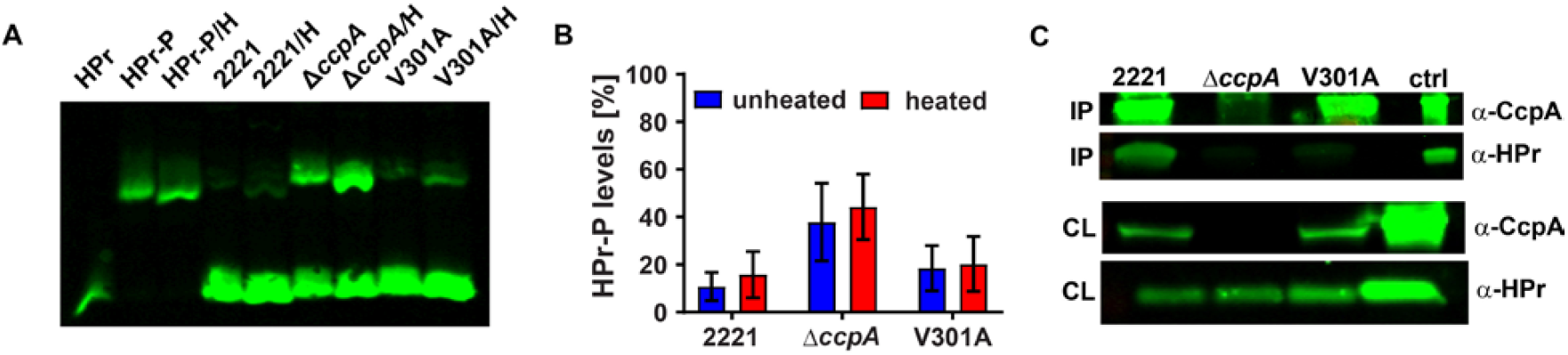
Characterization of the CcpAV301A mutant protein. (A) Representative Phostag Western blot (n=2) and (B) graphical representation of the levels of phosphorylated HPr in lysates of wild type (2221), *ccpA* deletion mutant (Δ*ccpA*) and the CcpAV301A mutant (V301A) grown to mid-exponential phase. Lanes marked with “H” indicate heated samples. Purified recombinant HPr and HPr∼P was used as controls. Error bars indicate standard deviation. (C) Co-immunoprecipitation of HPr from lysates of indicated strains using anti-CcpA antibody. “IP” and “CL” indicates immunoprecipitated material and cell lysate respectively. The antibody for each sub-panel is indicated on the right. Purified recombinant CcpA and HPr proteins were used as controls (ctrl).

To further test our hypothesis that CcpAV301A was not interacting with HPr in vivo, we performed immunoprecipitation reactions using anti-CcpA antibody which detects both the wild type and CcpAV301A mutant protein (Figure 2C, first and third panels). Strains MGAS2221, 2221Δ*ccpA*, and 2221-CcpA-V301A were crosslinked and harvested at mid-exponential phase. CcpA-containing complexes were immunoprecipitated using a polyclonal anti-CcpA antibody and then analyzed for the presence of HPr by western blotting using anti-HPr antibody (Figure 2C). By quantitative analysis, 10 times more HPr was immunoprecipated by CcpA antibody from strain MGAS2221 compared to strain 2221-CcpA-V301A. No HPr was immunoprecipated by CcpA antibody in strain 2221Δ*ccpA*. Taken together, we conclude that the V301A amino acid change essentially abolishes the ability of CcpA to interact with HPr both in vitro and in vivo.

### Inability of CcpA to interact with HPr impacts the CcpA transcriptome

Given that CcpA-HPrSer46∼P interaction is thought to be critical for CcpA activity (Deutscher *et al*., 2005, Deutscher *et al*., 2006, Deutscher, 2008, Gorke & Stulke, 2008, Stulke & Hillen, 2000), we next sought to test the hypothesis that the CcpAV301A change would result in a transcriptome similar to that observed for a complete CcpA knockout. To this end, we performed RNAseq analyses, in quadruplicate, for strains MGAS2221, 2221Δ*ccpA*, and 2221-CcpA-V301A grown to mid-exponential phase in THY. We did not observe any significant changes in *ccpA* transcript levels between MGAS2221 and 2221-CcpA-V301A, confirming that the V301A change does not affect transcription of *ccpA* itself (Supplementary Figure S1). Principal component analysis showed that the data were reproducible and, contrary to our hypothesis, that the transcriptomes of the three strains were quite distinct (Figure 3A). We assigned transcript levels as being significantly different when there was a mean difference of at least 1.5 fold and a P < 0.05 after accounting for multiple comparisons. We first sought to assess how the transcriptome of MGAS2221differed from 2221Δ*ccpA*. Deletion of *ccpA* resulted in significantly different transcript levels of 514 genes representing ∼31% of the 1667 genes in the MGAS2221 genome with sufficient transcript levels for analysis (Supplementary Table 1). More genes (361) had increased transcript levels in strain 2221Δ*ccpA* (i.e. were CcpA repressed) compared to genes (153) whose transcript levels were lower in 2221Δ*ccpA* (i.e. were CcpA activated), which is in accordance with previous observations (Carvalho *et al*., 2011, DebRoy *et al*., 2016, Kietzman & Caparon, 2011, Shelburne *et al*., 2008). The transcript levels of twelve virulence factor encoding genes were impacted by CcpA inactivation including *speB, speA2, sic, spd, ska, grab*, the *sag* operon, the *nga*-*slo* operon, and *prtS* (Table 2). Consistent with the central role of CcpA in carbon source acquisition and utilization, the transcript levels of genes encoding 13/14 of the phosphotransferase systems (PTS) present in MGAS2221 were significantly altered by CcpA inactivation as were genes encoding four ATP binding cassette (ABC) carbohydrate transporters (Table 3). Cluster of orthologous group (COG) analysis revealed that genes whose transcript levels were significantly impacted by CcpA inactivation were more likely to be in groups C (energy production and conversion) and G (carbohydrate transport and metabolism) relative to unaffected genes (Figure 3B).

**Table 2.** Influence of CcpA-HPr interaction on virulence gene regulation and DNA binding.

| M5005<br>spy# | Gene | CcpA<br>repressed<br>vs<br>activated | HPr<br>dependent vs.<br>independent | If HPr<br>dependent :<br>semi or fully | Bound<br>by<br>CcpA | Bound by<br>CcpAV301A |
| --- | --- | --- | --- | --- | --- | --- |
| 0139-0141 | *nga-slo* | Repressed | Dependent (*nga*) | Semi | No | No |
| 0341 | *prtS* | Repressed | Independent |  | Yes | Yes |
| 0351 | *spyA* | Repressed | Independent |  | No | No |
| 0562-70 | Streptolysin S operon | Repressed | Dependent | Semi | Yes | Yes |
| 0996 | *speA2* | Repressed | Independent |  | No | No |
| 1106 | *grab* | Activated | Dependent | Semi | No | No |
| 1540 | *endoS* | Activated | Dependent | Semi | No | No |
| 1684 | *ska* | Repressed | Independent |  | No | No |
| 1711 | *lmb* | Repressed | Independent |  | No | No |
| 1718 | *sic* | Repressed | Independent |  | No | No |
| 1735 | *speB* | Activated | Independent |  | Yes | Yes |
| 1738 | *spd* | Repressed | Independent |  | No | No |

**Table 3.** Influence of CcpA-HPr interaction on regulation and DNA binding of genes encoding carbohydrate transport systems of MGAS2221.

| **M5005 spy#** | **Putative transported carbohydrate** | **Transporter type<sup>a</sup>** | **CcpA repressed vs activated** | **HPr dependent vs. independent** | **Bound by CcpA** | **Bound by CcpAV301A** |
| --- | --- | --- | --- | --- | --- | --- |
| 0212-0219 | Sialic acid | ABC | Repressed | Independent | Yes | No |
| 0475 | B-glucoside | PTS | Repressed | Semi-dependent | No | No |
| 0521 | N-acetylglucosamine | PTS | Repressed | Independent | No | No |
| 0662 | Fructose | PTS | Repressed | Fully - dependent | No | No |
| 0780-0783 | Mannose/fructose | PTS | Repressed | Semi-dependent | No | No |
| 1058-1060 | Maltose | ABC | Repressed | Semi-dependent | Yes | Yes |
| 1067-1062 | Maltodextrin | ABC | Repressed | Semi-dependent | Yes | Yes |
| 1083-1079 | Cellobiose | PTS | Repressed | Semi-dependent | Yes | Yes |
| 1310-1308 | Unknown sugar | ABC | Repressed | Semi- dependent | No | No |
| 1399-1401 | Galactose | PTS | Repressed | Semi-dependent | No | No |
| 1479-1481 | Mannose | PTS | Repressed | Semi-dependent | Yes | Yes |
| 1542 | Sucrose | PTS | Activated | Semi-dependent | No | No |
| 1634-1633 | Lactose | PTS | Activated | Fully - dependent | No | No |
| 1664-1662 | Mannitol | PTS | Repressed | Semi-dependent | No | No |
| 1692 | Maltose | PTS | Repressed | Independent | Yes | Yes |
| 1746-1744 | Cellobiose | PTS | Repressed | Semi-dependent | Yes | No |
| 1784 | Trehalose | PTS | Repressed | Semi-Dependent | No | No |

<sup>a</sup>PTS - phosphotransferase system; ABC - ATP binding cassette

**Figure 3.**
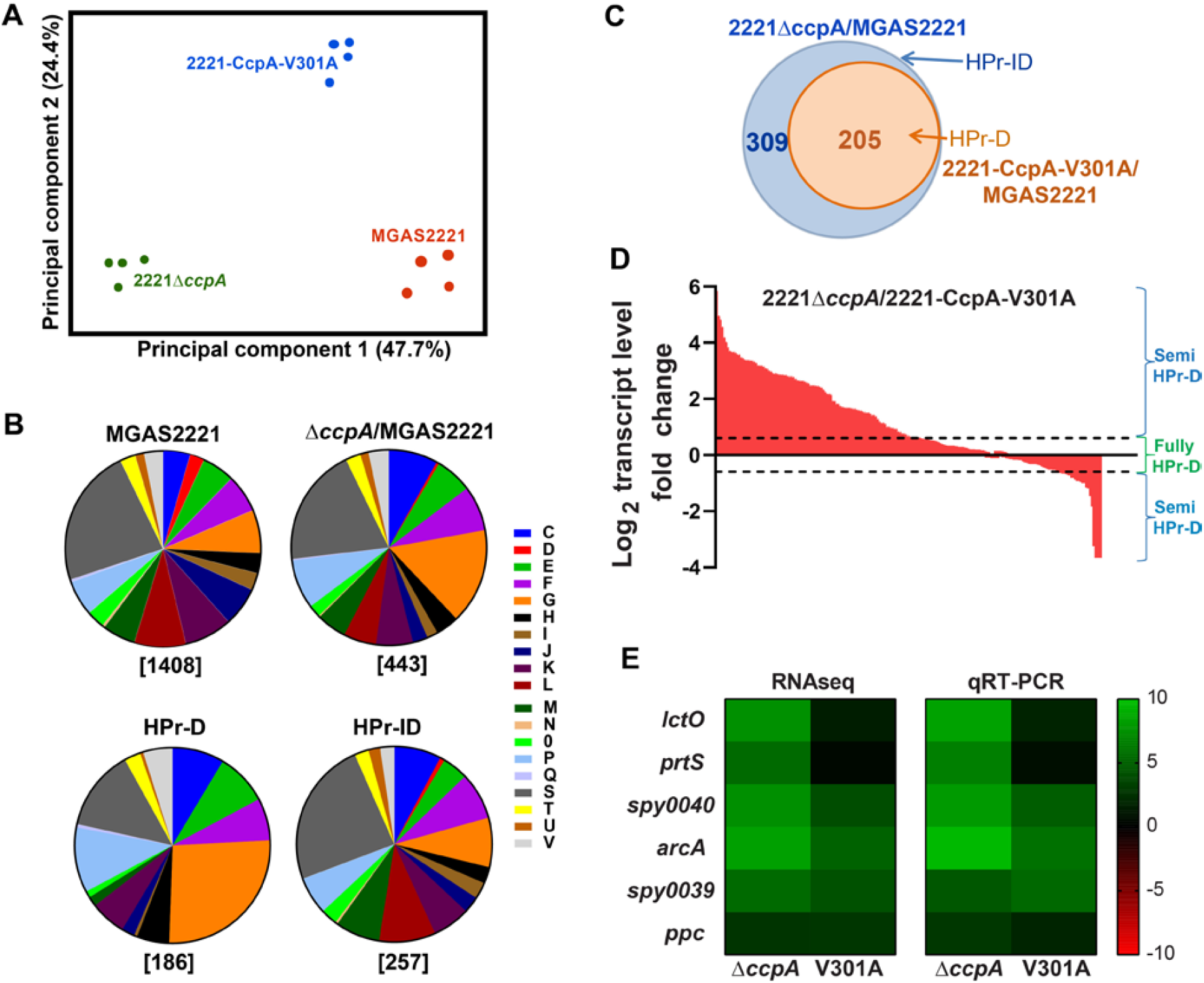
Impact of CcpAV301A mutation on the GAS transcriptome. (A) Principal component analysis showing that the transcriptomes of MGAS2221, 2221Δ*ccpA* and 2221-CcpA-V301A are distinct. Each strain has four biological replicates. (B) COG-based distribution of all genes in MGAS2221, genes impacted by *ccpA* deletion (Δ*ccpA*/MGAS2221) and the HPr-dependent (HPr-D) and HPr-independent (HPr-ID) genes. [C] Energy production and conversion; [D] Cell cycle control, cell division, chromosome partitioning; [E] Amino acid transport and metabolism; [F] Nucleotide transport and metabolism; [G] Carbohydrate transport and metabolism; [H] Coenzyme transport and metabolism; [I] Lipid transport and metabolism; [J] Translation, ribosomal structure and biogenesis; [K] Transcription; [L] Replication, recombination and repair; [M] Cell wall/membrane/envelope biogenesis; [N] Cell motility; [O] Posttranslational modification, protein turnover, chaperones; [P] Inorganic ion transport and metabolism; [Q] Secondary metabolites biosynthesis, transport, and catabolism; [S] Function unknown; [T] Signal transduction mechanisms; [U] Intracellular trafficking, se*cre*tion, and vesicular transport and [V].Defense mechanisms. (C) Venn diagram showing the subsets of *ccpA*-affected genes that are HPr-independent (HPr-ID) and HPr-dependent (HPr-D). The strain comparisons used to generate these subsets are indicated in their respective colors. (D) Waterfall plot showing the gradation in the magnitude of the transcriptional impact of the CcpAV301A mutation on the HPr-dependent genes. The fold change for HPr-dependent genes between strains 2221Δ*ccpA* and 2221-CcpA-V301A, as observed in the RNAseq, is plotted. The genes within the dotted line are fully HPr-dependent, while those outside are HPr semi-dependent. (E) Gene transcript levels of selected CcpA-impacted genes that exhibit varying transcriptional effects of the CcpAV301A mutation by RNAseq analysis were validated by targeted Taqman qRT-PCR analysis. Transcript levels in strains 2221Δ*ccpA* and 2221-CcpA-V301A are shown relative to wild type MGAS2221. For Taqman qRT-PCR, data are mean ± standard deviation of two biological replicates, with two technical replicates, done on two separate days (n = 8).

Next, we analyzed transcript levels in strain 2221-CcpA-V301 for genes whose transcript levels were significantly different between strains MGAS2221 and 2221Δ*ccpA*. If the CcpA-HPr interaction is critical to regulation of a particular CcpA-impacted gene then we would expect that transcript levels for said gene to be significantly different between strains MGAS221 and 2221-CcpA-V301A. However, of the 514 genes whose transcript levels were significantly affected by CcpA inactivation, only 205 (40%) also had significantly different transcript levels in strain 2221-CcpA-V301A relative to strain MGAS2221 (Figure 3C). We considered these to be HPr-dependent genes because eliminating the interaction between CcpA and Hpr impacted their transcript levels (Supplementary Table 1). The remaining 309 (60%) genes whose transcript levels were impacted by CcpA inactivation did not have significantly different transcript levels between strains MGAS2221 and 2221-CcpA-V301A and were therefore denoted as HPr-independent (Figure 3C) (Supplementary Table 1). A significantly higher percentage of the HPr-dependent genes were CcpA repressed (157/205, 77%) compared to HPr-independent genes (204/309, 66%, P = 0.01 by Fisher’s exact test). For the twelve virulence factor encoding genes/operons impacted by CcpA inactivation, only four were HPr-dependent (*nga*-*slo, sag* operon, *grab*, and *endoS*). Conversely, of the seventeen CcpA-regulated genes/operons encoding carbohydrate transport systems, fourteen were HPr-dependent (Table 2, 3). Similarly, COG analysis showed that, the HPr-dependent genes were more likely to be in category G (carbohydrate transport and metabolism) and less likely to be in category L (replication and repair), M (cell wall/membrane/envelop biogenesis), and S (function unknown) relative to HPr-independent genes (Figure 3B).

We next sought to determine whether the loss of CcpA-HPr interaction was equivalent to complete CcpA inactivation in terms of the absolute effect on gene expression by using fold-change (FC) comparison of the HPr-dependent genes. If the CcpA-HPr complex is essential for CcpA-regulation of a particular gene, then there should be no significant difference in transcript levels between strains 2221Δ*ccpA* and 2221-CcpA-V301A. In fact, similar transcript levels in these two strains were observed for only 78 (38%) of the 205 HPr-dependent genes, which we subsequently considered as fully HPr-dependent (i.e. the effect of eliminating CcpA-HPr interaction was equivalent to the effect of total CcpA inactivation) (Supplementary Table 1). For the remaining 127/205 (62%) genes, transcript levels were significantly different between strains 2221Δ*ccpA* and 2221-CcpA-V301A. Given that CcpA is primarily a repressor, for these 127 genes the loss of interaction with HPr resulted in transcript levels that were generally lower than complete CcpA inactivation, but the magnitude of effect varied for individual genes (Figure 3D). We designated these genes as semi-HPr-dependent (Supplementary Table 1). All four HPr-dependent virulence factor encoding genes/operons (*nga*-*slo* operon, streptolyin S operon, *grab*, and *endoS*) were semi-HPr-dependent (Table 2). Of the 14 HPr-dependent carbohydrate utilization encoding operons, only two (lactose and fructose transporters) were fully HPr-dependent (Table 3). We chose CcpA-regulated genes that exhibit varying degrees of transcriptional impact upon disrupting the CcpA-HPr interaction and verified the changes in transcript levels using targeted qRT-PCR (Figure 3E). Taken together, our data show that CcpA influences a significant proportion of the GAS genome even without HPr interaction and that the quantitative impact of HPr on CcpA-mediated regulation is distinct for different genes.

### In vivo interaction of CcpA and CcpAV301A with GAS chromatin

There are limited genome wide assessments of CcpA binding to DNA in pathogenic bacteria and none for a strain in which CcpA-HPr interaction has been abrogated (Willenborg *et al*., 2014). Therefore, we performed ChIP-Seq analysis for strains MGAS2221 and 2221-CcpA-V301A grown to mid-exponential phase in THY media to analyze CcpA chromatin occupancy profiles and assess how the reduced ability of CcpA to interact with HPr affects CcpA-DNA binding. The 2221Δ*ccpA* strain was used as a control to ensure the specificity of our findings.

We identified 76 CcpA binding sites in strain MGAS2221, including in the promoter region of *ccpA* itself which is consistent with a previous in vitro analysis of CcpA binding (Almengor *et al*., 2007) (Table 4). The genome-wide distribution of these sites is shown in Figure 4A. The lack of significant enrichment at any of these sites in strain 2221Δ*ccpA* indicates that these are true in vivo binding sites for the CcpA protein. The CcpA binding sites were mostly (67%) in predicted promoter regions (i.e. within 500 bps of a translational start site) (Fujita, 2009), which is in accord with other CcpA ChIP-Seq analyses (Antunes *et al*., 2012, Buescher *et al*., 2012, Willenborg *et al*., 2014). Of the 76 enriched regions, 46 sites (61%) were associated with genes whose transcript levels were significantly altered in strain 2221Δ*ccpA* relative to strain MGAS221. These 46 sites were associated with 45 operons containing 107 genes. The discrepancy between numbers of binding sites and operons affected is due to the occasional identification of multiple distinct binding sites for the same gene. Thus, only 107 (21%) of the 514 genes whose transcript levels were significantly impacted by CcpA inactivation in strain MGA2221were identified as being directly regulated by CcpA (Figure 4B). The vast majority of sites enriched in the ChIPseq analysis (88%) correlated with genes which had increased transcript levels in strain 2221Δ*ccpA* relative to MGAS2221 (i.e. were CcpA-repressed). By COG analysis, genes directly regulated genes by CcpA were significantly more likely to be in category C (energy production and conversion) and G (carbohydrate transport and metabolism) (Figure 4C).

**Table 4.**
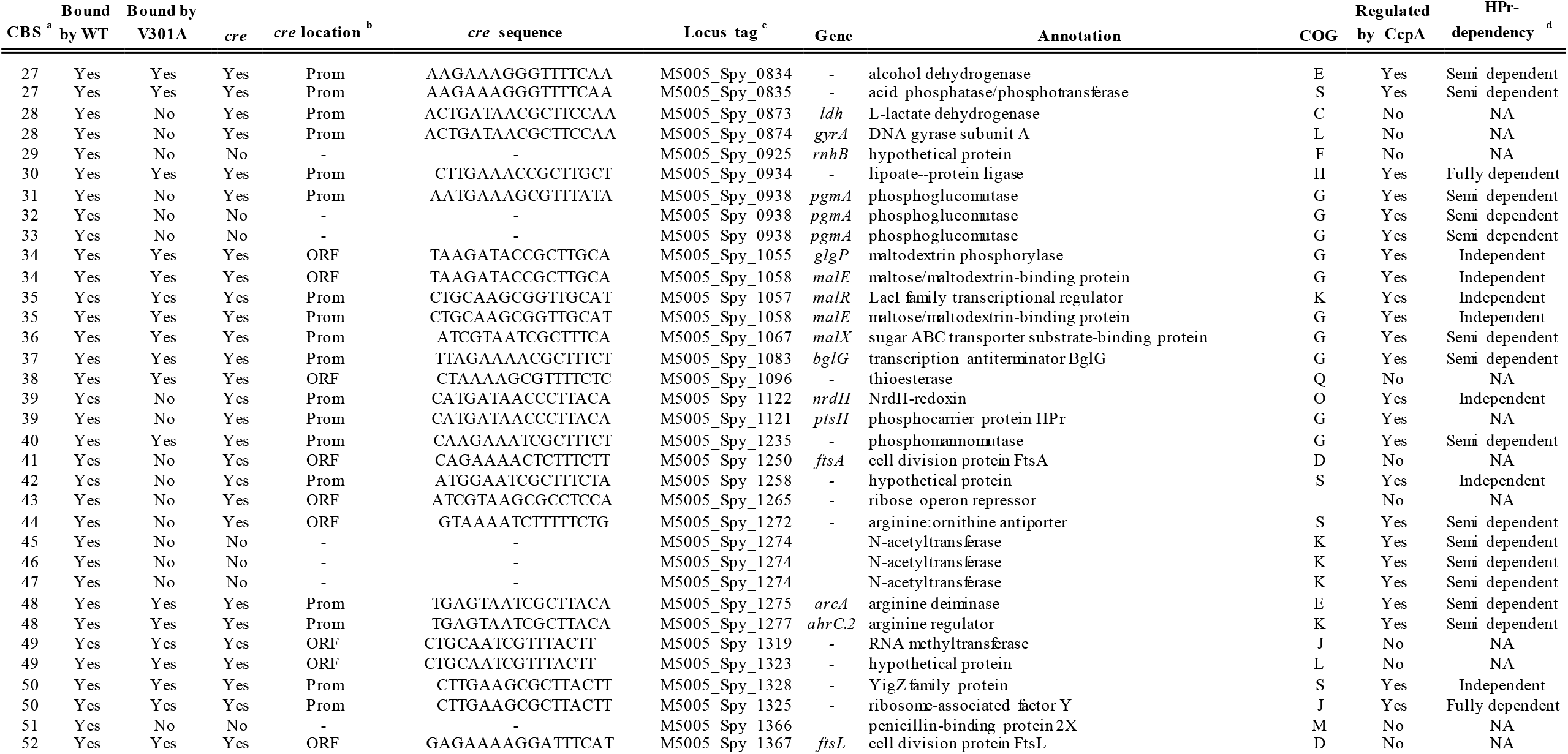
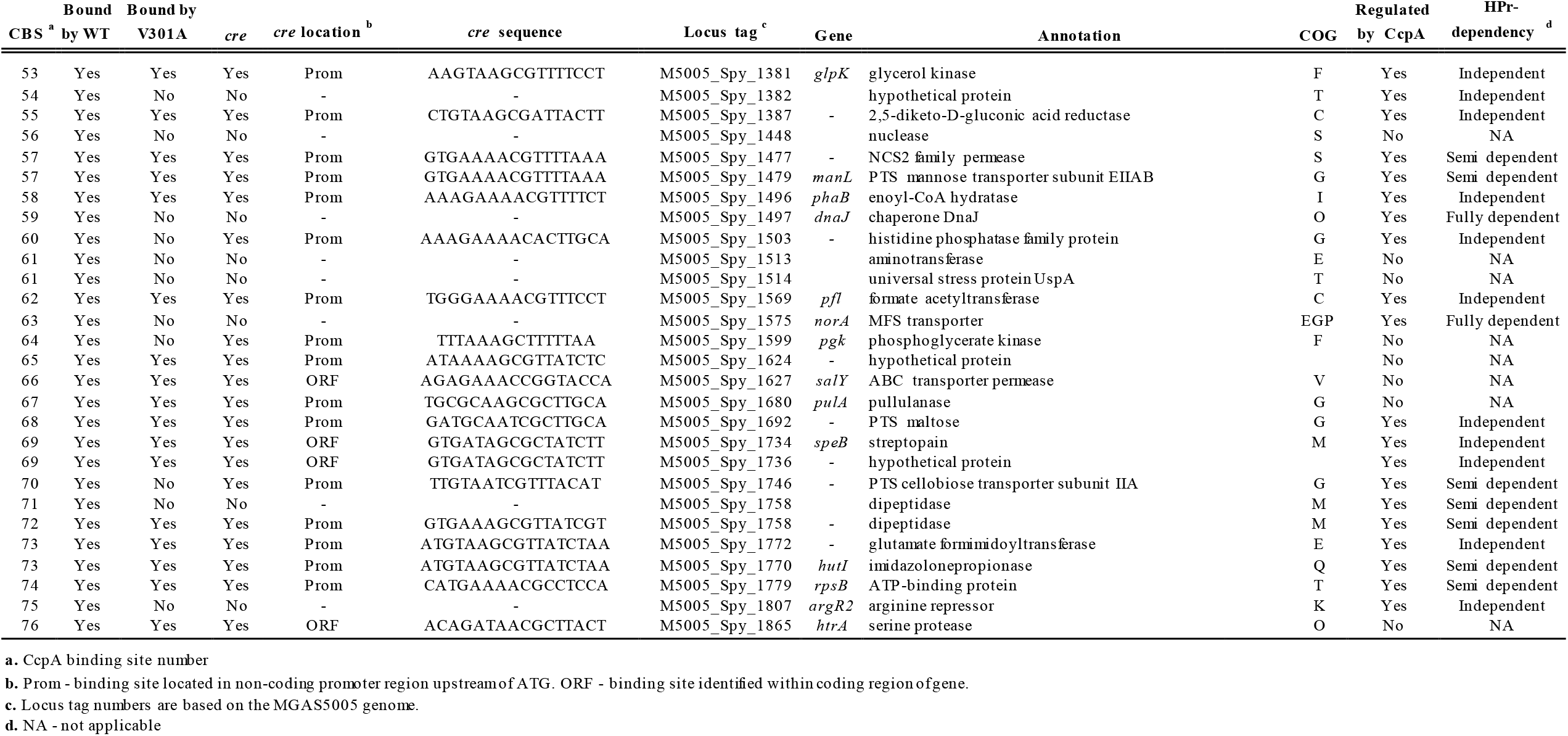
CcpA binding sites identified by ChIPseq analysis and associated genes.

**Figure 4.**
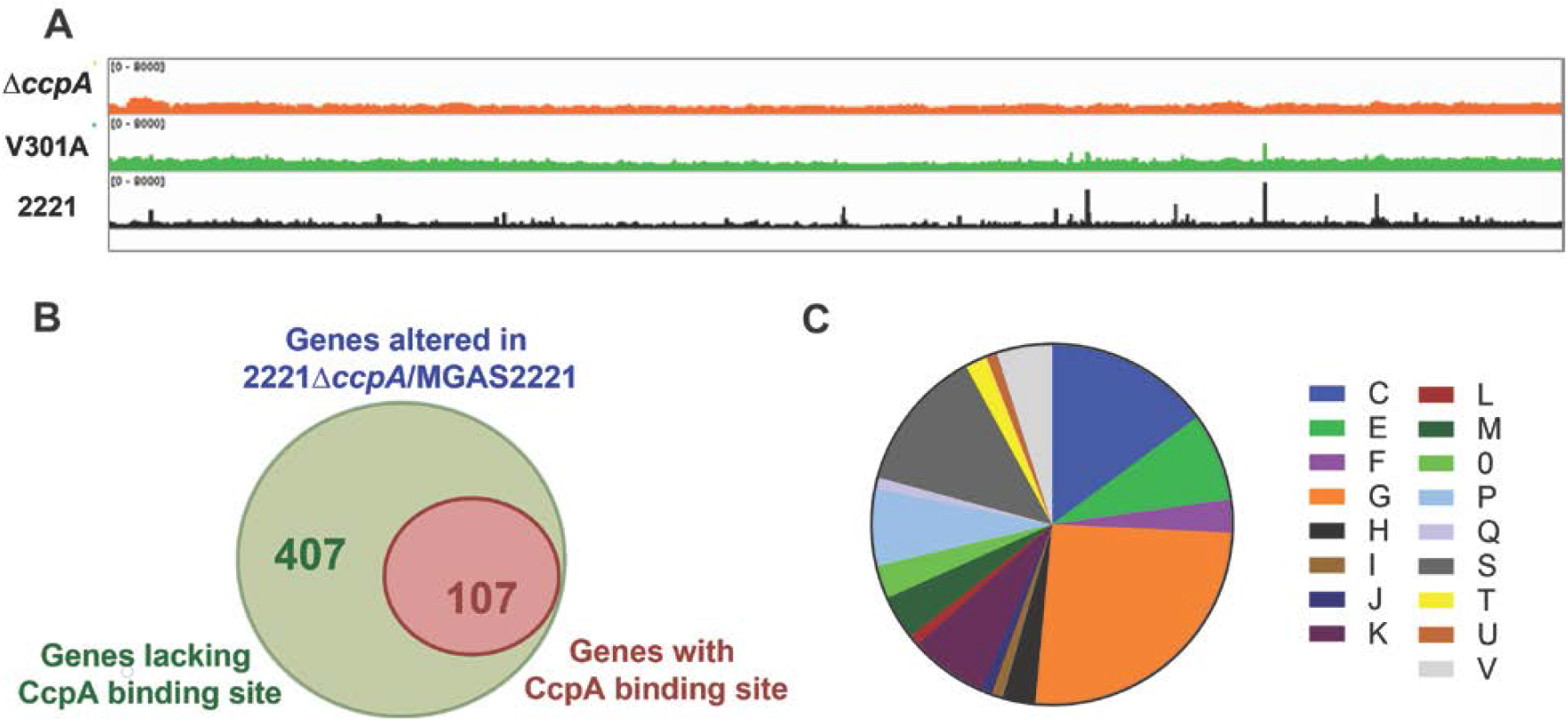
Characterization of in vivo DNA binding of CcpA. (A) Linear representation of indicated strains showing the CcpA binding sites as determined by ChIPseq analysis. (B) Venn diagram showing proportion of genes identified as CcpA-regulated by RNAseq that have CcpA binding sites as identified in our ChIPseq analysis. (C) COG distribution of genes that have CcpA binding sites and are transcriptionally altered in 2221Δ*ccpA*. Description of COG categories is provided in legend of Figure 3B.

Specific CcpA binding was identified for genes encoding four PTS systems known and putatively involved in the transport and utilization of maltose, cellobiose (two operons), and mannose, which stands in contrast to the 13 PTS gene/operons whose transcript levels were affected by CcpA inactivation. We also observed CcpA binding sites in genes/operons encoding three ABC carbohydrate transport systems for maltose, maltodextrin, and sialic acid. The central role of maltose/maltodextrin utilization in GAS pathophysiology is reflected by the five distinct CcpA binding sites for genes involved in the acquisition of these prevalent carbohydrates including that in the promoter region of the *pulA* gene encoding a cell-surface pullulanase important for both nutrient acquisition and adherence (Figure 5A). In addition to transporters, genes encoding glycerol kinase, phosphoglucomutase, and HPr, which are also involved in carbohydrate metabolism, had CcpA binding sites and were directly regulated by CcpA.

**Figure 5.**
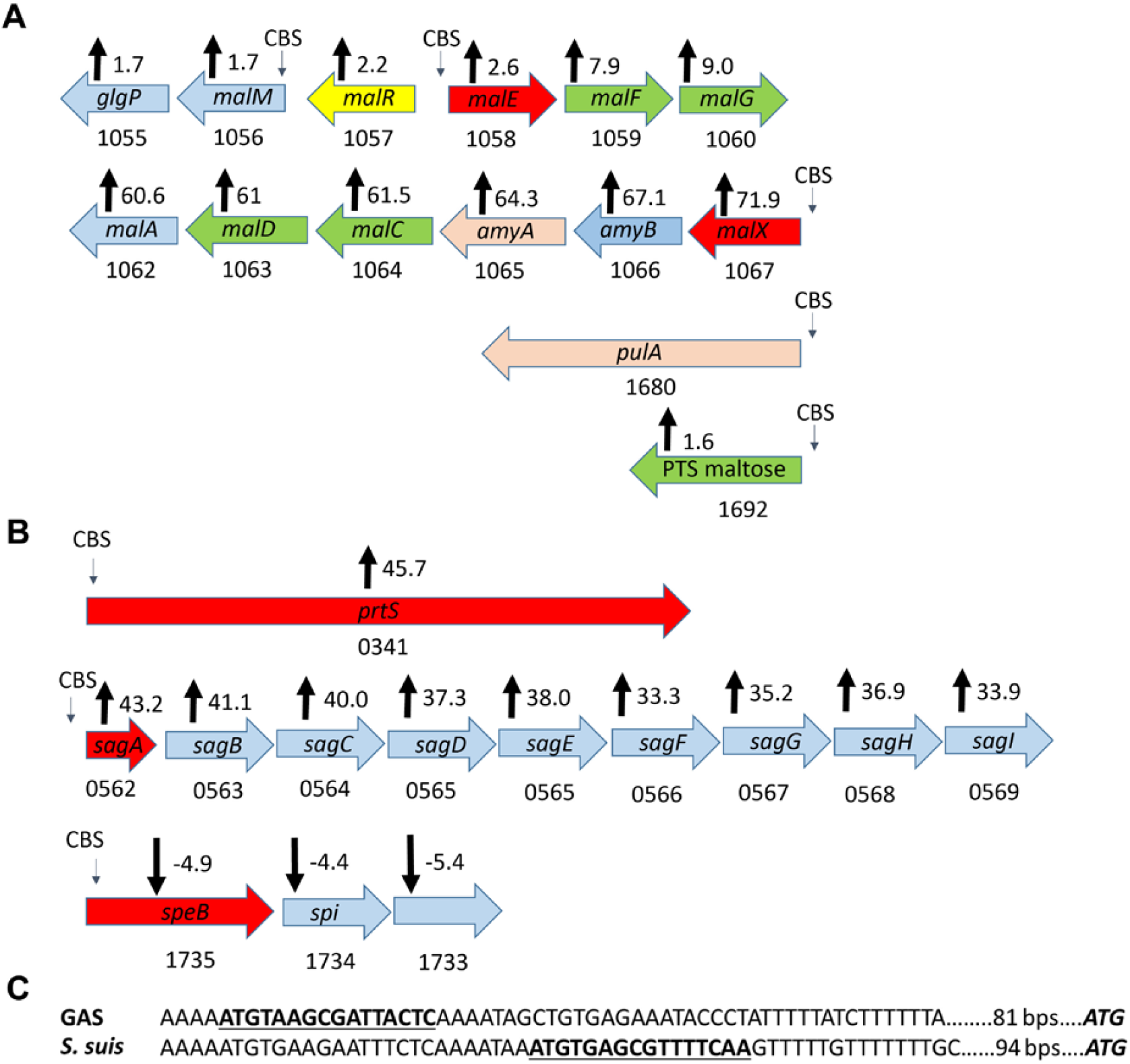
CcpA-bound sites in MGAS2221. Schematic representation of the binding sites for CcpA (CBS) in key (A) carbohydrate transport and (B) virulence factor encoding genes. Fold change in transcript levels upon CcpA inactivation is displayed with black arrows. Positive numbers indicate higher transcript levels in 2221Δ*ccpA* compared to MGAS2221. Genes in the schematic diagram are color coded to indicate function: blue – intracellular carbohydrate processing protein; yellow – transcriptional regulator; red – substrate binding lipoprotein; green – transport protein and beige – cell surface/se*cre*ted protein. (C) Sequence variation in the promoter region of the arginine deiminase (*arcA*) gene in GAS and *S. suis*. Enriched site for GAS and putative CcpA binding site for *S. suis* are highlighted.

CcpA enriched sites were also identified in the promoters of genes encoding various amino acid uptake and utilization pathways such as those involved in arginine, histidine, and glutamate metabolism. Although a recent CcpA study in *Streptococcus suis* did not identify in vivo CcpA binding for *arcA*, the initial gene in the arginine catabolism operon, *arcA* showed the strongest CcpA-mediated DNA enrichment for the entire MGAS2221 dataset. A comparison of the *arcA* promoters showed that whereas a CcpA binding site is present in GAS and predicted for *S. suis*, these sites are quite variable both in terms of composition and location, which may explain the divergent results (Figure 5C).

Importantly, DNA enrichment was observed for the promoter and/or 5’ coding regions of three key GAS virulence factor encoding genes (Figure 5B). Namely we observed in vivo binding of CcpA for *sagA*, the first gene in the operon encoding the key cytotoxin Streptolysin S, *speB*, which encodes an actively se*cre*ted broad-spectrum cysteine proteinase, and *prtS* which encodes an IL-8 degrading enzyme. No CcpA binding or effect on gene transcript level was observed for *cfa* (M5005_spy0981) which was previously reported to be directly regulated by CcpA (Kietzman & Caparon, 2009). Similarly, we did not observe enrichment for the other nine virulence factor encoding genes whose transcript levels were varied by CcpA inactivation (Table 2) suggesting an indirect mechanism for CcpA impact on these genes.

Given the large numbers of genes impacted by CcpA inactivation relative to the number of identified CcpA binding sites, it was reasonable to suspect that many of the genes might be secondarily affected by regulators under CcpA control. Indeed, RNAseq analysis showed that the transcript levels of 21 genes/operons encoding known and putative transcriptional regulators, including four two-component gene regulatory systems (TCS), were significantly affected by CcpA inactivation (Supplementary Table 1). Of these, however, we identified direct CcpA binding for only three genes encoding the transcriptional regulators BglG, ArgR2 and Spy1495 (Table 4). While the *bglG* and *argR2* genes had a CcpA-binding site in their promoters, *spy1495* was in an operon where the first gene (*spy1496*) was CcpA-bound. As previously noted, there was also CcpA enrichment in the intergenic region between *malE* and *malR* (Figure 5A), which encodes a maltose regulatory protein. Hence, *malR* could also be directly regulated by CcpA although the location of the binding site suggests that CcpA is more likely to influence *malE* transcription. There was no DNA enrichment for any TCS or the master regulator Mga (<u>m</u>ulti <u>g</u>ene <u>a</u>ctivator), which was identified to be directly regulated by CcpA in a previous study (Almengor *et al*., 2007). Given that none of the regulators directly controlled by CcpA are known or expected to have large transcriptomes (Shelburne *et al*., 2011), these data suggest that the broad impact of CcpA inactivation on GAS gene transcript levels is unlikely to be primarily mediated by regulators directly under CcpA control.

### Analysis of CcpAV301A in vivo DNA Binding

Of the 76 DNA loci bound by CcpA, 37 enriched regions which encompass 75 genes (12% of the CcpA transcriptome) were also enriched in strain 2221-CcpA-V301A (Figure 4A). All of the enriched sites in 2221-CcpA-V301 were associated with genes that also evidenced enriched sites in MGAS2221. All three of the virulence factor encoding genes bound by CcpA in strain MG22221, *prtS, sagA*, and *speB*, also evidenced binding in strain 2221-CcpA-V301A (Table 2 and 4). Similarly, five of the seven PTS/ABC carbohydrate transport encoding genes/operons (Table 3 and 4) bound by CcpA were also bound by CcpAV301A as was *arcA* (Table 5). In contrast, *spy1746*, which encodes the first gene in a cellobiose PTS operon, and *spy0212*, which encodes the first gene in the sialic acid uptake ABC operon were not bound by CcpAV301A. We also did not observe binding of the CcpAV301A protein to the promoter of either *ccpA, ptsH* (HPr) or the operon encoding the ATP synthase genes (Table 4). When considering all DNA binding sites for the CcpA protein in strain MGAS2221, those sites that were also bound in strain 2221-CcpA-V301A were more likely to have significantly different transcript levels following CcpA inactivation (P < 0.001 by by Fisher’s exact test).

### Characterization of CcpA binding sites not associated with differentially expressed genes

For 29 CcpA binding sites, we did not observe a significant difference in transcript levels for a nearby gene following CcpA inactivation. The majority of these binding sites were located in the middle of genes suggesting that the observed CcpA binding is likely non-functional. For nine genes, the DNA enrichment overlapped with promoter regions including four genes whose function would suggest regulation by CcpA (Table 4). These four genes were *pgi*, which encodes a glucose utilization protein, *plr*, which encodes an aldehyde dehydrogenase, *ldh*, which encodes an enzyme that converts lactate to pyurvate, and the previously mentioned *pulA*. Of these 29 CcpA enriched sites that were not associated with transcript level variation after CcpA inactivation, 11 were also enriched in strain 2221-CcpA-V301 including three genes encoding carbohydrate utilization proteins (*pgi, plr*, and *pulA*) as well as one site in a CRISPR operon and the *salY* gene, which encodes an ABC transport protein involved in lantibiotic se*cre*tion (Table 4). The combination of CcpA binding site location and putative function suggests that these genes may be directly regulated by CcpA under conditions distinct from those studied herein.

### Identification of CcpA binding motif

Using the ChIPseq data, we identified a single consensus motif in the enriched sites for MGAS2221 (Figure 6B) which was highly consistent with previously identified *cre* motifs from *Bacillus* species (Marciniak *et al*., 2012, Schumacher *et al*., 2004). This motif was present in 57 of the 76 CcpA enriched sites (75%). For the remaining 19 sites, no particular consensus motif was identified. We did not identify the *cre*2 sequence of *S. suis* or the flexible binding site of *Clostridium acetobutylicum* that were recently described (Willenborg *et al*., 2014, Yang *et al*., 2017). Compared to enriched areas with *cre* sites, enriched areas lacking *cre* sites were less likely to be in genes/gene promoters whose transcript levels were significantly affected by CcpA inactivation (P = 0.01 by Fisher’s exact test), suggesting that the *cre*-containing enriched DNA sites were more biologically impactful under the studied conditions.

**Figure 6.**
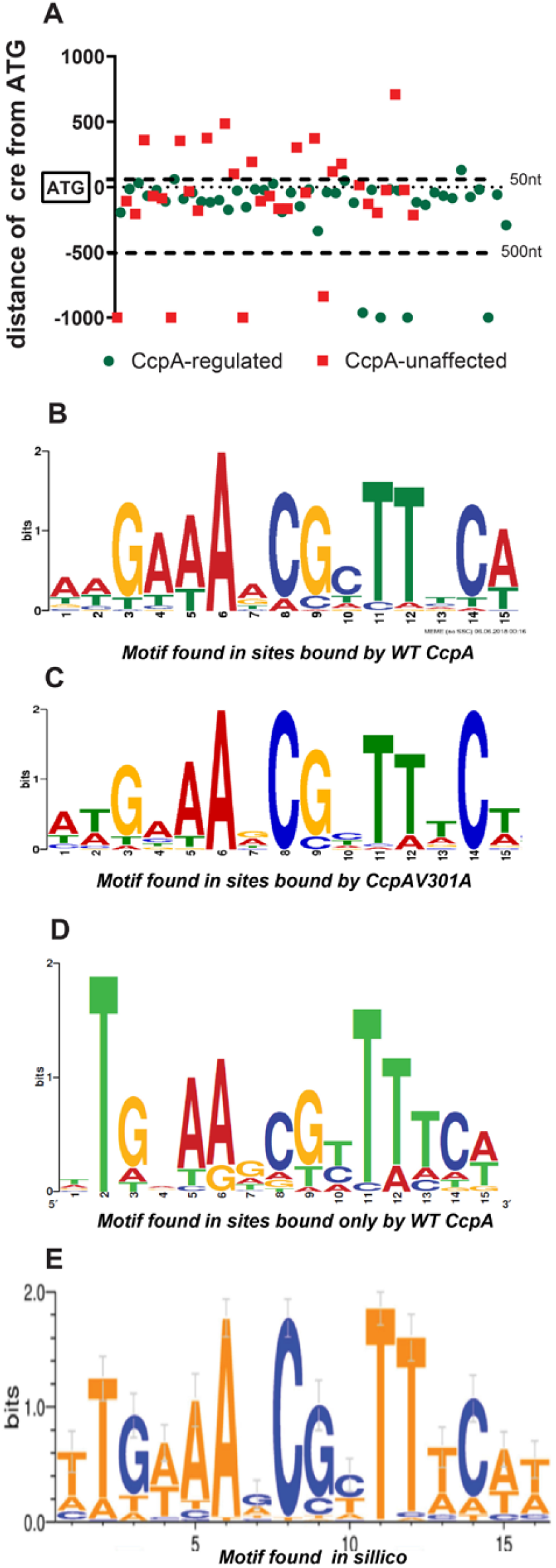
Consensus motifs identified in CcpA binding sites. (A) Scatter plot showing the distance from the translational start site (TSS) of enriched sites bound by the wild type CcpA protein that contain a consensus motif. Genes associated with the enriched sites are color coded to indicate whether they are (CcpA-regulated) or not (CcpA-unaffected) transcriptionally impacted upon CcpA inactivation. Enriched sites that are located farther than 1000nt from the TSS are not plotted. Weblogo representation of the consensus *cre* motif identified from CcpA binding sites in strains (B) MGAS2221 and (C) 2221-CcpA-V301A. (D) The *cre* motif found in sites that were bound by the wild type CcpA but not the CcpAV301A mutant. (E) The consensus motif identified in our previous study by in sillico analysis of CcpA-regulated genes in three different GAS serotypes (DebRoy *et al*., 2016).

The *cre* sites that were associated with genes whose transcript levels were affected by CcpA inactivation, were significantly more likely to be located within 500nt upstream and 50nt downstream of the start codon compared to *cre* sites that were near genes not impacted by CcpA inactivation (P <0.001 by Fisher’s exact test) (Figure 6A). Relative to our previously described in silico derived *cre* (Figure 6E, (DebRoy *et al*., 2016)), the major difference for our ChIPseq derived *cre* was a lack of predominance of T at positions 1 and 2 (Figure 6B). We compared the *cre* sites that we found using the ChIP method in this study to those that we had previously predicted in MGAS2221 using in silico analysis of RNAseq data (DebRoy *et al*., 2016). Both studies were conducted on mid-exponential cells grown in THY. We had predicted 72 *cre* sites in silico compared to the 57 we found by ChIPseq in this study. Of these sites, only 29 were in common with 20 being transcriptionally altered upon CcpA inactivation in both studies.

Importantly, the ChIPseq analysis enabled identification of 14 additional binding sites that are near genes that have altered transcript levels in a Δ*ccpA* mutant but were not predicted by the in silico approach. These include the *cre* sites identified in the promoter of *prtS, sagA* and the three *cre* sites in the PTS and ABC transport systems for maltose/maltodextrin. Conversely, the in silico method predicted 23 DNA loci that were transcriptionally impacted by CcpA inactivation in our study but were not bound by CcpA in our ChIPseq analysis. These genes included those encoding the transcriptional regulator FruR, an alcohol dehydrogenase, and the ABC transporter Spy1310.

When considering sites enriched in 2221-CcpA-V301A only (Figure 6C), we identified a motif that was highly similar to that for strain MGAS2221. The occurrence of an A at position 15 was reduced in the *cre* consensus derived from DNA loci in strain 2221-CcpA-V301A compared to that derived from MGAS2221. Of 57 *cre* sites in MGAS2221, 37 were bound in strain 2221-CcpA-V301, and we did not identify any enriched *cre* containing sites that were present in strain 2221-CcpA-V301 but not in MGAS2221. For *cre* sites that were bound only in strain MGAS2221 but not in the strain 2221-CcpA-V301A (Figure 6D), the occurrence of a T at position 2 was absolute and markedly increased relative to all *cre* sites in MGAS2221. Thus, the ChIPseq data show that although GAS *cre* sites have a similar architecture to other Gram-positive bacteria, in silico prediction of GAS CcpA binding using motifs is problematic.

### Impact of CcpAV301A mutation on magnitude of transcript level variation and degree of DNA binding

Given our finding of the differential transcript level impact of the CcpAV301A change (Figure 2D), we sought to further study how interruption of CcpA-HPr interaction impacted CcpA function by assessing whether the CcpAV301A amino acid variation quantitatively impacted CcpA gene regulation and DNA binding. For clarity of analysis, we only examined genes whose transcript levels were significantly increased by CcpA inactivation in MGAS2221 (e.g. were CcpA repressed), that evidenced in vivo DNA binding by CcpA, and were monocistronic or the first gene in an operon. We identified 33 genes that met these criteria, all but one of which (*argR2*) was associated with a *cre* (Supplementary Table 2). Using these genes, we tested the hypothesis that the CcpAV301A polymorphism would result in less release of catabolite repression compared with total CcpA inactivation. Indeed, for 27/33 genes, transcript levels were significantly higher in strain 2221Δ*ccpA* relative to 2221-CcpA-V301A, up to 57-fold but with a broad range (Supplementary Table 2).

Next, we sought to determine whether this differential impact on transcript levels correlated with a change in DNA binding affinity as determined by amount of DNA immunoprecipitated using anti-CcpA antibody. We used SYBR qPCR to quantify precipitated DNA from the promoter regions of six genes that evidenced CcpA binding for both MGAS2221 and 2221-V301A-CcpA strains by ChIPseq. For all six genes, we observed significantly more DNA precipitation in MGAS2221 compared to 2221-CcpA-V301A and compared to strains 2221Δ*ccpA* (Figure 7A). However, we did not identify a consistent relationship between quantitative differences in amount of DNA precipitated and gene transcript level variation observed in our RNAseq results (Figure 7B). For example, there was markedly more DNA from the *ppc* and *lplA* promoters precipitated in strain MGAS2221 compared to 2221-CcpA-V301A (Figure 7A), yet there was no significant difference in *ppc* and *lplA* gene transcript levels between strains 2221 and 2221-CcpA-V301A relative to 2221Δ*ccpA* (Figure 7B). Conversely, there was only a modest difference in the amount of *malX* promoter DNA precipitated from MGAS2221 compared to 2221-CcpA-V301A (Figure 7A), yet we observed marked variation in transcript levels between MGAS2221 and 2221-CcpA-V301A relative to 2221Δ*ccpA* (Figure 7B). Taken together, we conclude that abrogation of CcpA-HPr interaction quantitatively impacts CcpA function both in terms of the effect of CcpA on gene transcript levels and DNA binding affinity, but decreases in CcpA affinity for promoter DNA due to the V301A alteration does not strictly correlate with a subsequent impact on gene transcript level under the studied conditions.

**Figure 7.**
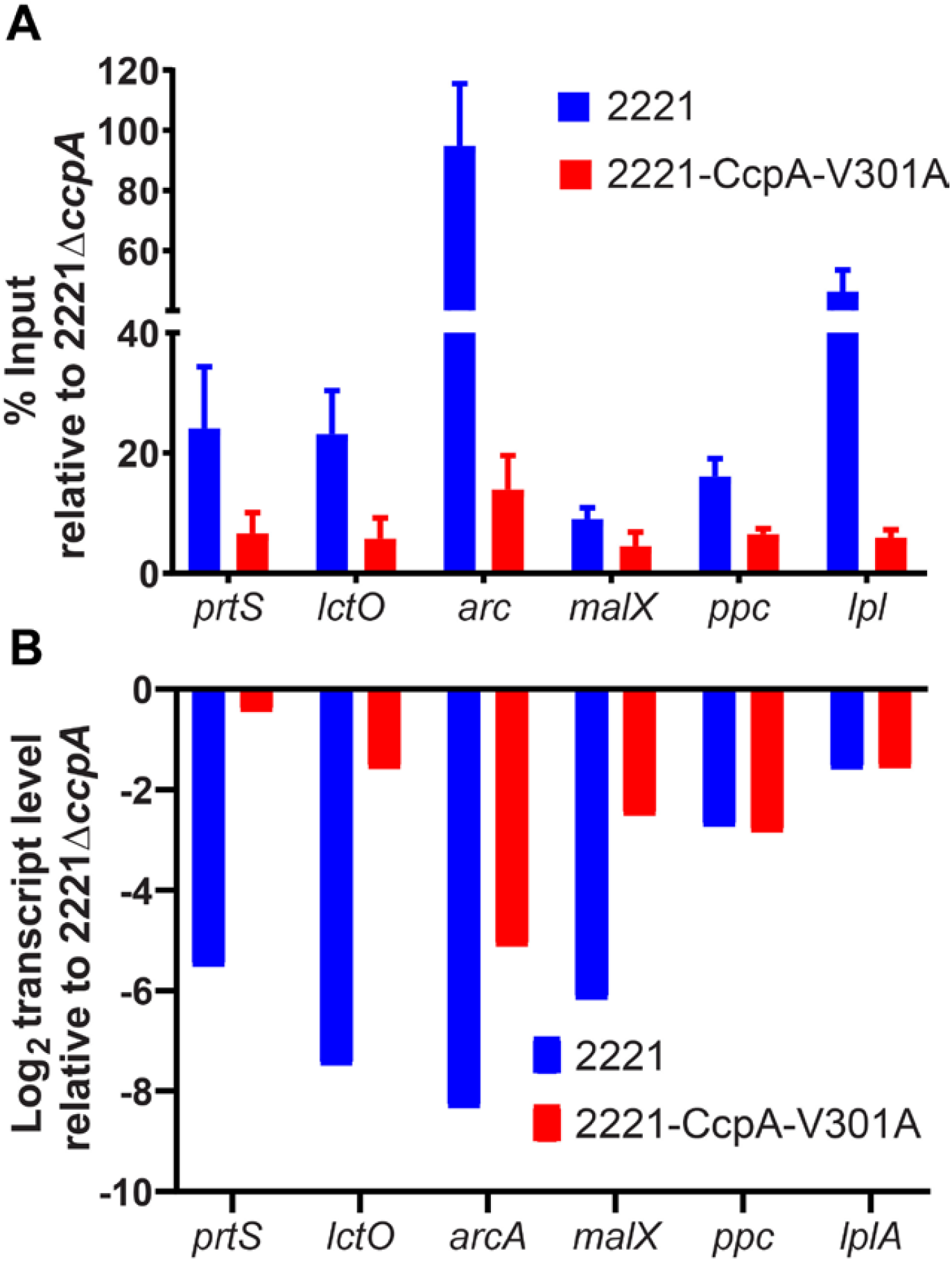
Quantitative impact of CcpAV301A alteration on gene transcript level and DNA binding. (A) SYBR qRT PCR analysis of DNA precipitated using anti-CcpA antibody from indicated strains (legend inset) for specific promoters (indicated on x-axis). (B) Transcript level changes as reported in our RNAseq data for the six genes in panel A.

## Discussion

The complex interplay between basic metabolic processes and bacterial pathophysiology is being increasingly appreciated (Eisenreich *et al*., 2010, Pacheco *et al*., 2012, Rohmer *et al*., 2011). For Gram-positive bacteria, CcpA stands squarely at this interface given its central role in controlling preferred carbon utilization pathways and regulating virulence factors in a diverse array of human pathogens (Iyer *et al*., 2005, Mendez *et al*., 2012, Shelburne *et al*., 2008, Vega *et al*., 2016). The primary link of CcpA to the energy status of the bacterial cell is via its interaction with HPr, but study of HPr in many important Gram-positive bacteria has been limited due to its essential nature (Fleming *et al*., 2015, Willenborg *et al*., 2014). Herein we sought to extend our knowledge about how CcpA links metabolic and virulence processes through a comprehensive identification of in vivo CcpA binding sites in the major human pathogen group A *Streptococcus*. Additionally, we created a strain in which CcpA cannot interact with HPr, to delineate HPr-dependent and HPr-independent aspects of CcpA function. Our findings show that CcpA directly regulates the genes encoding several critical GAS virulence factors and that >50% of genes in the GAS CcpA transcriptome are impacted by CcpA independent of HPr.

A key contribution of our study is the first genome wide analysis of in vivo CcpA DNA binding for a β-hemolytic streptococci. We identified 76 CcpA binding sites in GAS, the specificity of which were demonstrated by the lack of enrichment in a CcpA knockout strain. The only other available CcpA ChIPseq analysis in streptococci was performed in *Streptococcus suis* (Willenborg *et al*., 2014) and identified 58 DNA loci bound by CcpA at the mid-exponential phase of growth as was studied herein. Some loci, such as those in the promoters of *glpK, malX*, and *pgmA* were identified in both studies, while others like *arcA, manLMN, pgi* and *eno* were not. In MGAS2221 we found the *arcA* site to be the most strongly enriched peak and the most derepressed gene upon *ccpA* deletion. In contrast, the *S. suis* study did not identify enrichment of the *arcA* or the *manLMN* promoters even though these genes were impacted by CcpA inactivation. Additionally, we did not identify the *cre*2 motif present in the *S. suis* dataset, perhaps because that motif was identified at the stationary phase of growth, which we did not investigate.

A common observation of both this and the *S. suis* study (Willenborg *et al*., 2014), and of CcpA investigations in other pathogens (Antunes *et al*., 2012), is that direct CcpA binding accounts for only a small percentage of genes affected by CcpA inactivation. For example, of the genes encoding the 14 known carbohydrate PTS systems in MGAS2221, the transcript levels of 13 were affected by CcpA inactivation, but only four are directly regulated. This narrow direct impact of CcpA on the PTS systems was also observed in *S. suis*, where CcpA directly impacted only two of the 14 PTS systems present. The *manLMN* operon in GAS and *Streptococcus pneumoniae* has been shown to impact a diverse array of other carbohydrate transporters (Abranches *et al*., 2003, Abranches *et al*., 2006, Fleming & Camilli, 2016, Vadeboncoeur & Pelletier, 1997). We hererin established that in GAS, CcpA binds the promoter of the *manLMN* operon in vivo (Table 3) and thus could indirectly account for some of the broad impact of CcpA inactivation on carbohydrate transport systems despite only directly binding a small number of genes encoding these systems. While it has been postulated that the broad CcpA transcriptome may be due to the impact of CcpA on other regulators (Antunes *et al*., 2012, Carvalho *et al*., 2011, DebRoy *et al*., 2016, Seidl *et al*., 2009), we found that CcpA only bound the promoters of a small number of other regulator encoding genes (Table 4) which is not consistent with a broad effect of CcpA on the GAS regulatory network. The marked increase in HPr∼P observed following CcpA inactivation may account for a substantial proportion of the indirect effects of CcpA given the known role of HPr∼P in a broad array of regulatory processes (Deutscher *et al*., 2005).

Another key finding of this work was our identification that CcpA directly binds to genes encoding three critical GAS virulence factors, *sagA, prtS*, and *speB*. How CcpA mechanistically impacts the expression of the *sag* operon, and subsequent production of the key cytotoxin streptolysin S, has been an object of study for the past 12 years since the original observation that CcpA inactivation strikingly increases steptolysin S production (Kinkel & McIver, 2008, Shelburne *et al*., 2008). Our group as well as others have identified in vitro binding of CcpA to the *sagA* promoter (Kinkel & McIver, 2008, Shelburne *et al*., 2008) whereas an in vivo study showed no precipitation of the *sagA* promoter using a CcpA antibody (Kietzman & Caparon, 2009). Although both MGAS2221 and HSC5, the strain used in the aforementioned study (Kietzman & Caparon, 2009), both contain the *sagA cre* site we identified, the area upstream of the *cre* site is quite divergent for the two strains which could explain the differential in vivo binding results. It has been shown that the *manLMN* operon and the β-glucoside PTS (Table 3), which we identified herein as being directly and indirectly regulated by CcpA respectively, impacts streptolysin production (Braza *et al*., 2020, Sundar *et al*., 2017) suggesting that CcpA may regulate streptolysin S production both directly and indirectly. In concert with our data, CcpA was identified as binding in vivo to the gene encoding suilysin, the pore-forming toxin of *S. suis*, (Willenborg *et al*., 2014) and to bind in vitro to the *sagA* promoter of *S. anginosus* (Bauer *et al*., 2018). We identified direct binding of CcpA to *speB*, in accordance with previous studies that have used DNA pulldown and fluorescence polarization methods to demonstrate this interaction (Kietzman & Caparon, 2009, Shelburne *et al*., 2010).

Interestingly, *speB* was one of the very few genes directly activated by CcpA. Additionally, our identification that CcpA directly regulates *prtS*, which encodes a critical IL-8 degrading enzyme, marks the first identification of in vivo binding of a regulator to this important GAS virulence factor. The evolutionary rationale for having such critical and diverse virulence factor encoding genes under direct control of a global metabolic regulator is not clear, but CcpA has also been shown to directly and indirectly impact the expression of key virulence factors in a range of Gram-positive bacteria from Clostridia to Staphylococci (Abranches *et al*., 2008, Chiang *et al*., 2011, Seidl *et al*., 2006, Varga *et al*., 2004).

Although HPr has long been recognized as critical to CcpA function, study of HPr in major human pathogens such as staphylococci and streptococci has been limited due to its essential nature, including the critical Ser46 residue (Fleming *et al*., 2015, Willenborg *et al*., 2014). We also were unable to affect HPrSer46∼P through a mutagenesis approach and thus sought to interrupt CcpA-HPr∼P interaction by modifying the critical CcpAV301 residue. Surprisingly, although the V301A mutation abrogated CcpA-HPr interaction both in vitro and in vivo, more than half of the genes in the CcpA transcriptome still demonstrated CcpA-based repression in strain 2221-CcpA-V301A, indicating the capacity of CcpA to function independently of HPrSer46∼P in GAS. We were able to correlate the preserved function of the CcpAV301A protein through ChIPSeq analysis which revealed continued enrichment for numerous DNA promoters in strain 2221-CcpA-V301A. Our identification of CcpA functioning independent of HPr echoes findings from a recent ChIPseq study in *S. suis* which found that CcpA can bind to promoters of and regulate several genes in both exponential and stationary phase of growth (Willenborg *et al*., 2014). Given the absence of HPrSer46∼P in the stationary phase, they postulated that the CcpA binding and regulation observed in the stationary phase probably occurred independent of HPr (Willenborg *et al*., 2014). These findings stand in contrast to those from *Staphylococcus xylolus* in which HPrSer46∼P is absolutely essential for CcpA-mediated catabolite repression (Jankovic & Bruckner, 2002). Our findings are more in concert with studies in *B. subtilis*, the non-pathogenic gram positive model bacterium, in which the impact of a HPrS46A mutation is essentially an overall dampening of gene regulation in terms of magnitude (Lorca *et al*., 2005). Indeed, our ChIPseq analysis revealed that enrichment of promoter DNA was typically reduced in strain 2221-CcpA-V301A compared to the wild-type even when significant changes in transcript levels were identified, suggesting that the CcpAV301A bound to *cre* sites with lower affinity. Similarly, we found that the magnitude of effect on gene transcript levels due to the inability of the CcpAV301A mutant protein to bind HPrSer46∼P varied such that genes could be grouped into classes that are strongly, mildly or not impacted by the abrogation of CcpA-HPr interaction.

One possible purpose of the varied magnitude of effects on gene transcript levels observed in the 2221-CcpA-V301A strain would be to facilitate a broad array of transcriptional responses to carbohydrate availability and intracellular energy status (Paluscio *et al*., 2018). GAS infects numerous different infection sites such as skin, oropharynx, blood and muscle (Cole *et al*., 2011, Cunningham, 2000, Walker *et al*., 2014). Each of these sites have unique nutritional and environmental makeup. The dependence/independence of CcpA from HPr∼P could assist with fine-tuning regulation of pertinent genes by CcpA such that site specific resources can be utilized optimally to aid the pathogen. For example, a recent study compared the GAS transcriptome during necrotizing fasciitis to growth in standard laboratory medium (Kachroo *et al*., 2020). Some of the most highly upregulated genes in Kachroo et. al,. including *sagA, arcA*, and the cellobiose PTS operon, were directly bound by CcpA in our study, all of which we classified as HPr-dependent. Conversely, the transcript levels of the directly CcpA bound, HPr-independent genes of the sialic acid operon, *prtS, ackA*, and *hpr* showed no significant upregulation during necrotizing fasciitis (Kachroo *et al*., 2020). These findings are consistent with the concept that CcpA can maintain repression of these genes even under metabolically unfavorable conditions when HPrSer46∼P levels would be expected to be low.

In conclusion, herein we present a genome wide analysis of chromatin occupancy by CcpA which reveals that CcpA directly regulates multiple key GAS virulence factors in addition to a broad array of critical metabolic genes. Through use of a CcpA isoform incapable of interacting with HPr, we have delineated not only the role of HPr in CcpA-mediated gene regulation but also the ability of CcpA to function at certain sites independently of its key co-factor. These findings extend the mechanistic understanding of how CcpA contributes to the pathophysiology of Gram-positive bacteria.

## Experimental Procedures

### Bacterial strains, media and growth conditions

All strains used in this study are listed in Table 1. Group A *Streptococcus* (GAS) strains were routinely grown in Todd-Hewitt media supplemented with 0.2% yeast extract (THY) at 37°C with 5%CO_2_. A single amino acid change in *ccpA* was engineered into the wild type MGAS2221 strain by homologous recombination using the pBBL740 plasmid as described previously (Horstmann *et al*., 2018), to *cre*ate the isoallelic strain 2221-CcpA-V301A.

### Recombinant proteins and antibodies

Site directed mutagenesis (primers in Supplementary Table 3) was used to introduce the V301A mutation into an existing clone of GAS *ccpA* (Shelburne *et al*., 2008) in the pET-His2 vector to generate the pET-His2-V301A plasmid, which was transformed into *E. coli* BL21/pLysS. Wild type CcpA and V301A mutant protein were induced overnight at 18°C, and purified to >95% homogeneity as described previously (Shelburne *et al*., 2010) and extensively buffer exchanged to 20mM Tris/HCl pH 7.5, 200mM NaCl. Recombinant GAS HPr was overexpressed, phosphorylated and purified as described previously (Shelburne *et al*., 2010). For the generation of polyclonal antibodies purified wild type CcpA and HPr proteins were used to immunize rabbits and affinity-purified antibody was obtained (Covance, Denver, PA).

### Protein-Protein interaction analysis

Surface Plasmon Resonanace (SPR) analysis was performed at 25°C using BIAcore T200 instrument (GE Healthcare). Wild type CcpA and V301A mutant proteins were immobilized on sensor chip CM5 via amine coupling. PBS (phosphate buffered saline (8.06 mM Na2HPO4 and 1.94 mM KH2PO4, pH 7.4, 2.7 mM KCl, 137 mM NaCl)) was used as running buffer for immobilization and binding experiment. Sensor chip surface was activated by injecting a 1:1 mixture of 0.4 M of 1-ethyl-3-(3-dimethylaminopropyl) carbodiimide hydrochloride and 0.1 M of N-hydroxysuccinimide over the flow cell surface at 5µl/min for 7 minutes. After the surface was activated, CcpA (10 µg/ml in 10mM NaOAc pH 5.0) and CcpAV301A (20 µg/ml in 10mM NaOAc pH 5.0) were injected over different flow cells.

Sensor surface was deactivated with an injection of 1.0 M ethanolamine-HCl pH 8.5 at 5µl/min for 7 minutes. Approximately, 3700 response units (RU) of CcpA and 4200 RU of CcpAV301A were immobilized. A reference flow cell was prepared with activation and deactivation steps but no protein was immobilized. Two-fold dilutions of HPr or HPrSer46∼P from 1.25 µM to 80 µM in PBS were flown over the immobilized proteins at 20 µl/min for 30 seconds. All SPR responses were baseline corrected by subtracting the response generated from the corresponding reference surface. Double-referenced SPR response curves (with the buffer blank run further subtracted) were used for affinity determination. The equilibrium response of each injection was collected and plotted against the concentration of injected protein. A one-site binding (hyperbola) model was fitted to the data (GraphPad Prism 4) to obtain the equilibrium dissociation constant *K*_D_.

### PhosTag gel

Recombinant HPr protein was phosphorylated and purified as described previously (Shelburne *et al*., 2010). Cell lysates of GAS strains were prepared, separated on 12.5% SuperSep Phostag gels and detected using a polyclonal anti-HPr antibody (Horstmann *et al*., 2014) as described previously (Horstmann *et al*., 2018). Experiments were repeated at least twice using samples collected on separate days.

### Co-immunnoprecipitation

Strains MGAS2221, 2221Δ*ccpA*, and 2221-CcpA-V301A were grown to mid-exponential phase in THY, crosslinked with EGS [ethylene glycol bis(succinic acid *N*-hydroxysuccinimide ester)] and formaldehyde and harvested. Pellets were sonicated and CcpA-containing complexes were immunoprecipitated using a polyclonal anti-CcpA antibody and then analyzed for the presence of HPr by western blotting using a polyclonal anti-HPr antibody and the Odyssey imaging system as described previously (Horstmann *et al*., 2015).

### Analysis of transcript levels

For RNA seq analysis, RNA was isolated from four replicate cultures for each strain grown to mid-exponential phase in THY (OD ∼ 0.5) using the RNeasy kit (Qiagen) and processed as previously described (Horstmann *et al*., 2014). RNAseq data analysis was performed as previously described (Sanson & Flores, 2020). As the MGAS2221 genome is not publicly available, we used the MGAS5005 genome as the reference genome, which is identical to MGAS2221 (Sumby *et al*., 2005) in gene content. For Taqman real-time qRT-PCR, strains were grown in duplicate on two separate occasions to mid-exponential phase in THY and processed as described previously (Horstmann *et al*., 2014). The gene transcript levels between MGAS2221 and either 2221Δ *ccpA* or 2221-CcpA-V301A was compared using an ordinary one way ANOVA. Primers and probes used are listed in Supplementary Table 3.

### Chromatin Immunoprecipitation (ChIP) and sequencing (ChIPseq)

For ChIP analysis, MGAS2221, 2221Δ*ccpA* and 2221-CcpA-V301A strains were grown to mid-exponential phase in THY. Crosslinking was performed with 1% formaldehyde and quenched with 0.125M glycine. DNA was sheared using a Diagenode Bioruptor Plus and immunoprecipitated with anti-CcpA antibody. We performed ChIPseq for two independent replicates of each strain. ChIP sequencing was performed in the Advanced Technology Genomics Core (ATGC) at MD Anderson Cancer Center. Briefly, Illumina compatible Indexed libraries were prepared from 12ng of Diagenode Biorupter Pico sheared ChIP DNA using the KAPA Hyper Library Preparation Kit (Roche). Libraries were enriched with 2 cycles of PCR then assessed for size distribution using the 4200 TapeStation High Sensitivity D1000 S*cre*enTape (Agilent Technologies) and quantified using the Qubit dsDNA HS Assay Kit (Thermo Fisher). Equimolar quantities of the indexed libraries were multiplexed, 9 libraries per pool. The pool was quantified by qPCR using the KAPA Library Quantification Kit (Roche) then sequenced on the Illumina NextSeq500 sequencer using the 75nt high-output flow cell. Raw FASTQ files from DNA sequencing were trimmed through Trimmomatic v. 0.36 (Bolger *et al*., 2014). A sliding window quality trimming was performed, cutting once the average quality of a window of three bases fell <30. Reads shorter than 32 bp after trimming were discarded. Resulting data were aligned to Genbank reference genome of *Streptococcus pyogenes* MGAS5005 (NC_007297.2) using Bowtie2 v. 2.3.0 (Langmead & Salzberg, 2012), without allowing mismatch. Resulting SAM files were converted to BED format using sam2bed v. 2.4.33. For each sample, the coverage per base was determined using coverageBed v. 2.27.1 from BEDTools (Quinlan & Hall, 2010). Subsequent processing was done using in-house Perl scripts. For that, data were normalized to the same average read depth and mean coverage per base was determined for each pair of replicates (Sharma *et al*., 2017). To select peaks for CcpA in MGAS2221 and 2221-CcpA-V301A, we compared the coverage base per base of both conditions with 2221Δ*ccpA*, and selected those bases scoring ≥2.0 fold over 2221Δ*ccpA*. Final peaks were obtained by merging selected bases using mergeBed v. 2.27.1. The sequences of assembled peaks were retrieved and submitted to Multiple Em for Motif Elicitation v. 5.0.0 (MEME) (Bailey & Elkan, 1994, Bailey *et al*., 2009) for motif searching. The expected number of sites was set to one per sequence and the minimum motif width was set to 15 bp (Sharma *et al*., 2017). A single statistically significant motif (E-value <1e −12) was recovered for MGAS2221 and 2221-CcpA-V301A and these matched the known CcpA binding consensus (DebRoy *et al*., 2016). The E-value is derived by MEME, from the motif’s log likelihood ratio, taking the motif length and background DNA sequence into account. Manual visualization of identified motifs and closest coding genes (CDS) on *S. pyogenes* MGAS5005 genome was performed using Integrative Genomic Viewer v. 2.3 (IGV) (Thorvaldsdottir *et al*., 2013).

## Acknowledgements

This work was supported by RO1 AI089891 (S.A.S) from the National Institues of Health, CONICYT-PIA Program AFB170001 of the Center for Mathematical Modeling (M.L), FONDECYT N° 1190742 (M.L), Center for Genome Regulation FONDAP 15090007 (M.L), Mesa Minería - Consorcio de Universidades del Estado de Chile – CUECH (M.L) and Fondo de Innovación para la Competitividad - FIC2018 - 6^ta^ región, Gobierno Regional Chile (M.L).

We acknowledge the CCSG P30 CA016672 funds for the Bioinformatics Shared Resource and the Sequencing and Microarray facility at MD Anderson Cancer Center.

We would like to acknowledge Kunal Rai and Christopher Terranova for their knowledge and guidance in developing ChIPseq methodology for GAS.

## Author Contributions

S.D., S.A.S., V.A., G.G., M.L. and M.H. designed the research, S.D., S.A., N.H. and S.A.S. did the experiments, S.D., X.L., S.A.S., A.R.F, V.A., G.G., V.M., M.L. and M.H. performed data analysis and S.D., X.L., M.L., M.H and S.A.S. wrote the manuscript. All authors reviewed the manuscript.

## Competing Financial Interests

The authors have no competing financial interests.

## Notes

### Competing Interest Statement

The authors have declared no competing interest.

