## Supplementary Figure 1 for "Genome-wide analysis of in vivo CcpA binding with and without its key co-factor HPr in the major human pathogen group A *Streptococcus*"

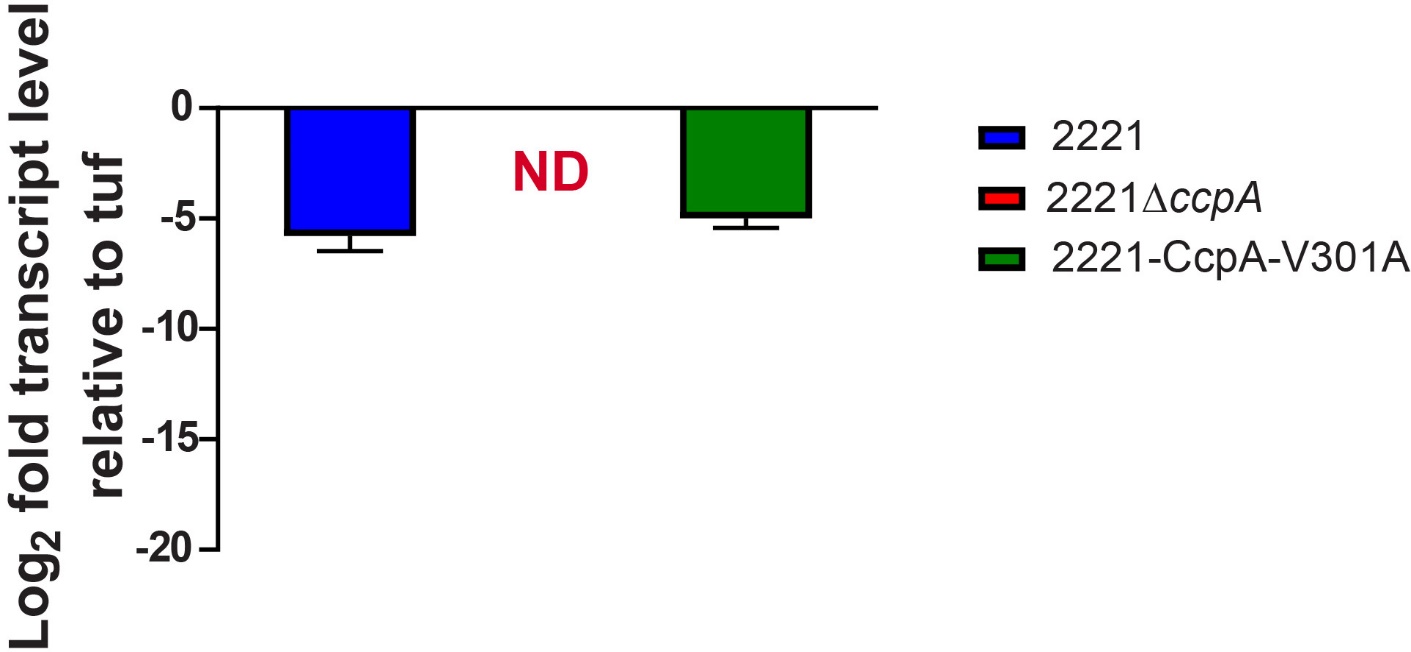

**Supplementary Figure 1. Transcript levels of *ccpA* in different GAS strains.** Gene transcript levels of *ccpA* were analysed by targeted Taqman qRT-PCR analysis in strains listed in the legend. Data are mean ± standard deviation of two biological replicates, with two technical replicates, done on two separate days (n = 8).
